# The scaffold protein CasL regulates T cell migration by restricting membrane blebbing

**DOI:** 10.1101/2024.12.12.628177

**Authors:** Liz A Kurtz, Hope E Shearer, Rosanne Trevail, Menelaos Symeonides, Mobin Karimi, Nathan H Roy

## Abstract

T cell migration into inflamed tissue is a key control point in the inflammatory response and relies on integrin interactions with their endothelial ligands. Here, we identify the signaling scaffold CasL (Hef1, NEDD9) as a central regulator of integrin-dependent migration in primary T cells. We found CasL is specifically needed for efficient migration on ICAM-1, but not VCAM-1 coated surfaces. While WT T cells migrating on ICAM-1 form an actin-rich cell front and move smoothly, T cells lacking CasL instead form numerous, aberrant membrane blebs. The abnormal blebbing observed in CasL KO T cells likely stems from diminished F-actin in the cell front coupled with increased contractile forces behind the nucleus, suggesting CasL regulates the cytoskeletal architecture in migrating T cells. Importantly, using an *in vivo* allogeneic hematopoietic transplant model we found that CasL promotes T cell migration into inflamed peripheral tissue, but was dispensable for trafficking to secondary lymphoid organs. Together, these results indicate CasL functions to control the balance of cytoskeletal components during integrin-dependent migration and highlight the importance of integrin signaling for proper migration into inflamed tissue.

## Introduction

The ability of leukocytes to migrate into and within peripheral tissue is a critical component of the inflammatory response and is required for adaptive immunity. However, unchecked immune infiltration can lead to tissue damage, chronic inflammation, and autoimmunity. A key control point of the inflammatory response is the process of transendothelial migration, in which T cells migrate out of the bloodstream into peripheral tissue by crossing the vascular endothelia ^1^. In response to inflammatory stimuli, local endothelial cells upregulate chemokines and adhesion molecules (such as selectins and integrin ligands) that initiate T cell rolling and adhesion to the endothelial wall, followed by cell spreading and ultimately migration across the barrier ^2^. Integrin interactions with endothelial ligands are critical for multiple steps in this process, including firm adhesion and migration. The primary T cell integrins involved in migration along and through the endothelia are LFA-1 (α_L_ß_2_) and VLA-4 (α_4_ß_1_) which bind their endothelial ligands ICAM-1/2/3 and VCAM-1, respectively ^1,2^. In addition to numerous *in vitro* and *in vivo* studies linking T cell integrins to transendothelial migration (for example,^3–6^), the importance of LFA-1 is underscored by the severe migration defects and resulting immunodeficiency in patients with Leukocyte Adhesion Deficiencies (LAD) I and III; genetic disorders in which patients lack the B2 integrin chain and thus have no LFA-1 (LAD-I), or lack an important adaptor and signaling protein downstream of LFA-1, kindlin-3 (LAD-III) ^7–12^.

Integrins clearly promote firm adhesion of T cells to the endothelia, however adhesion alone is not sufficient for successful T cell transmigration. To traverse the vascular wall and migrate within peripheral tissues, T cells must undergo substantial cytoskeletal reorganization. In general, the cytoskeletal architecture of migrating T cells is governed by the localized activity of Rho family GTPases and their downstream effector proteins, ultimately causing T cells to adopt a polarized morphology characterized by the generation of branched actin in the front of the cell driving outward protrusions and myosin-based contractility in the rear pulling the back of the cell forward (for example, ^13–17^). Importantly, T cell migration is strikingly plastic and can alternate between different migration modes depending on the physiological context ^18–20^. Therefore, the relative contributions of actin polymerization, myosin contraction, and adhesive contacts during T cell migration vary depending on the biochemical and biophysical properties of the environment and the cell signaling events that interpret and respond to those cues. During adhesive-based migration involving integrins, T cells display a broad actin-rich front and distinct uropod in the rear (for example ^13,21^). Interestingly, interacting with ICAM-1 coated surfaces alone triggers T cells to spread, polarize, and immediately migrate, showing the LFA-1/ICAM-1 interaction initiates all the necessary signaling and cytoskeletal changes needed for migration ^13,15,17,22–24^. In contrast, T cells migrating on the related integrin ligand VCAM-1 do not form a clear lamellipodium but instead form multiple protrusions and migrate in a more amoeboid fashion ^21,25^. The difference in migration on ICAM-1 and VCAM-1 can be easily distinguished when shear flow is applied, as T cells will migrate against the direction of shear on ICAM-1 but with the direction of shear on VCAM-1 ^21,24–28^. These studies suggest that the LFA-1/ICAM-1 interaction triggers a unique signaling and cytoskeletal response in migrating T cells. While many cytoskeletal regulatory proteins, such as Rho GTPases their effectors, have been identified to be important in this context (reviewed in ^29^), the signaling intermediates that link LFA-1 engagement to cytoskeletal responses are far less known.

One family of signaling scaffolds that have been implicated in integrin-based cell migration in a variety of cell types are the Cas family proteins. This family includes 4 members, p130Cas, CasL (HEF1, NEDD9), EFS/SIN, and HEPL (CASS4). All of the family members share a similar domain structure, with the salient feature being a large substrate binding domain that contains numerous tyrosine phosphorylation sites to coordinate adaptor protein binding and downstream signaling ^30^. Elegant studies on p130Cas have shown that this domain physically opens in an accordion-like fashion and can only be phosphorylated when force is applied across the protein, suggesting that at least this family member acts as a mechanosensor ^31,32^.

Functionally, Cas proteins are known to regulate integrin-dependent migration and cytoskeletal responses in non-hematopoietic cells ^33^. In particular, p130Cas and CasL have been shown to localize to integrin adhesion sites and coordinate downstream cytoskeletal remodeling (for example ^34–40^). In human cancers, hyperphosphorylation and/or expression changes of Cas proteins has been linked to more aggressive tumors and worse prognosis, further emphasizing their role in coordinating migration (reviewed in ^41^).

T cells primarily express the Cas family member CasL. Extensive biochemical work in immortalized T and B cell lines showed that CasL becomes heavily tyrosine-phosphorylated in response to T or B cell receptor stimulation and/or integrin ligation, and can form protein-protein interactions with cytoskeletal signaling proteins like FAK that are involved in migration ^42–48^. More recent functional studies using primary T cells from CasL knockout mice indicate that CasL is needed for a robust Ca^2+^ response after T cell receptor ligation, likely due to disrupted cytoskeletal-dependent TCR microcluster organization ^49^. Interestingly, both T and B cells isolated from CasL knockout mice display migration defects in an *in vitro* transwell migration assay, however, CasL knockout mice have a normal antibody response to vaccination, indicating these T and B cells are largely functional *in viv*o ^50^. These studies point toward a role for CasL in cytoskeletal responses and migration, although it is still unclear if, and in what context, CasL contributes to T cell function. Recently, we found that CasL becomes heavily phosphorylated when T cells are migrating on ICAM-1 coated surfaces, suggesting CasL may be a scaffold for integrin signaling in migrating T cells ^17^. Based on these findings, we hypothesized CasL may be particularly important for lymphocyte migration in inflammatory settings, where integrin-based migration is crucial to enter inflamed tissue. Therefore, we probed the role of CasL in integrin dependent migration both *in vitro* and *in vivo* with the goal of revealing new mechanisms that govern T cell migration and trafficking.

## Results

### CRISPR deletion of CasL in primary CD4^+^ T cells impairs migration

To determine if CasL plays a role in T cell migration we first generated primary, activated CD4^+^ T cells lacking CasL using a CRISPR/Cas9 system ^51^. Briefly, primary CD4^+^ T cells were isolated from Cas9 expressing mice, activated with anti-CD3 and anti-CD28, and then transduced with retroviral particles containing either non-targeting (NT) or specific guide RNAs to CasL (herein referred to as CasL KO) (**Fig 1A**). Importantly, this protocol allows for gene editing to occur after activation bypassing any potential effects of CasL knockout on T cell activation. Using this protocol, we were able to greatly reduce the protein levels of CasL in primary T cells as displayed in comparison to T cells from CasL germline knockout mice (herein referred to as CasL^-/-^) (**Fig 1B**). Importantly, this procedure did not affect T cell health or expansion after activation (**Fig 1C**). Previous studies indicated that T cells isolated from CasL knockout mice, as well as T cells knocked-down for CasL *ex vivo*, displayed defects in migration in a transwell setting ^50,52,53^, therefore we first assessed the migratory capabilities of CasL KO T cells using a similar transwell assay. Control (NT) and CasL KO T cells were serum starved then applied to transwell inserts with 3µm pores and allowed to migrate toward media containing 1% FBS in the bottom chamber for 2 hours. In comparison to control T cells, T cells lacking CasL had severe defects migrating through a transwell (**Fig 1D**). These data show that we can reliably generate primary T cells lacking CasL and these cells recapitulate the previously observed migration defects in a transwell migration assay.

**Figure 1:**
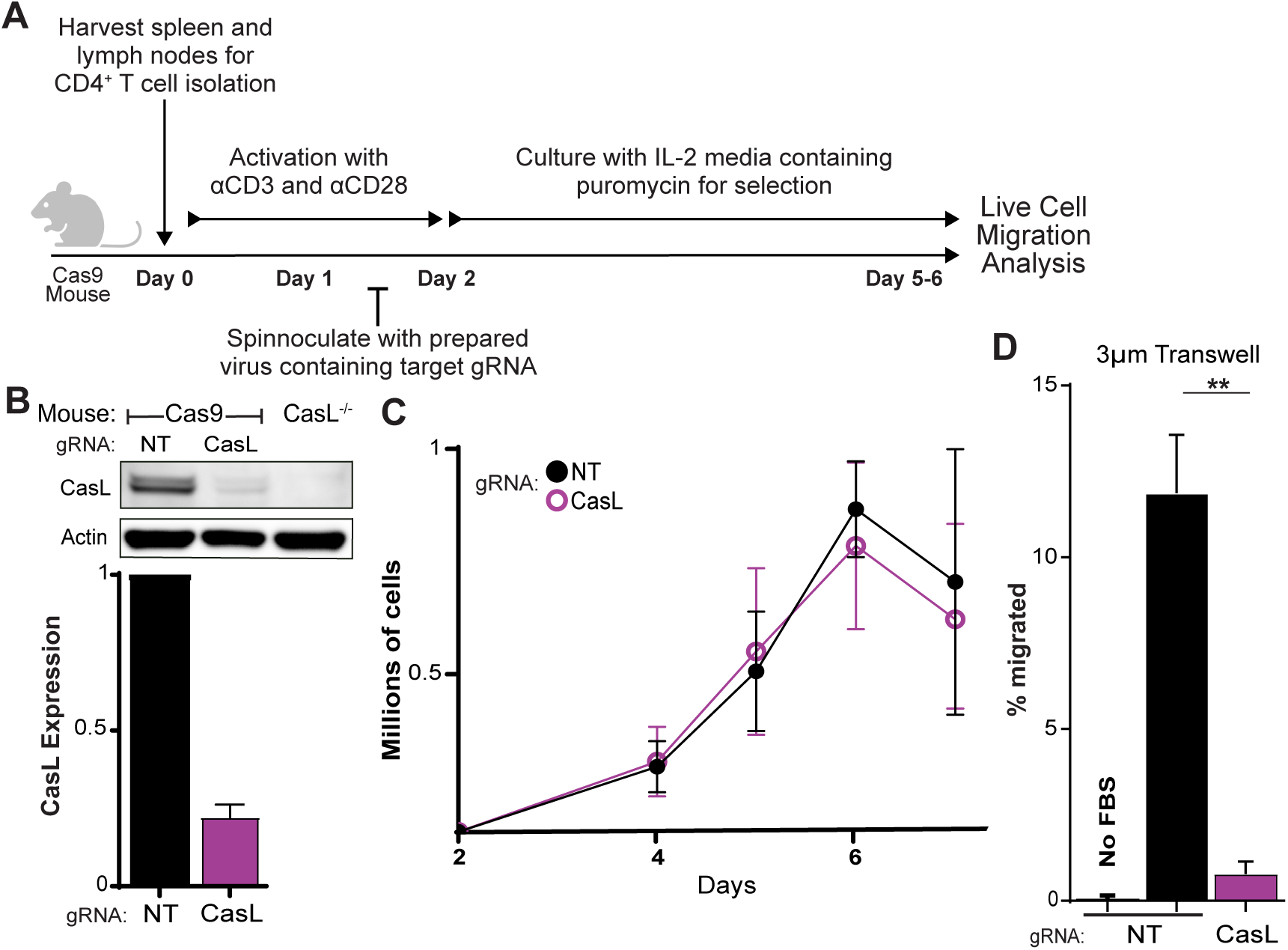
CRISPR Knockout of CasL in primary CD4+ T cells impairs migration. **(A)** Workflow to generate primary mouse CD4^+^ T cells lacking CasL. **(B)** Western blot of whole cell lysates from non-targeting gRNA transduced T cells (WT), CasL gRNA transduced T cells (CasL KO), and germline knockout CasL^-/-^ T cells, probed with anti-CasL. Quantification from n=8. **(C)** Growth curve comparing NT and CRISPR CasL KO cell number in culture on days 2-7 (n=3). **(D)** Percent migration of CD4^+^ T cells through a 3μm transwell toward 1% FBS over the course of 2h (n=3). T-test, ** = p < 0.01

### CasL is required for normal migration on ICAM-1 but not VCAM-1 coated surfaces

T cell integrins are crucial for multiple aspects of T cell motility, including traversing endothelial barriers and patrolling within peripheral tissue (as reviewed in ^2,20^). To determine if CasL is needed for T cell migration on integrin ligands, we allowed control or CasL KO T cells to migrate on ICAM-1 or VCAM-1 coated chamber slides and imaged them using time-lapse microscopy at low magnification for 10 minutes. Individual migrating cells were identified and manually tracked using ImageJ. Migrating cells were defined as cells that moved more than 2 cell lengths and stayed in the field of view for the entire imaging period. Cells passing this criteria were considered motile and included in downstream analysis of migratory parameters (**Supp Fig 1**). Plotting cell tracks from a representative experiment revealed that control T cells had longer, more directional tracks compared to CasL KO T cells when migrating on ICAM-1 coated surfaces. (**Fig 2A**). Interestingly, migration on VCAM-1 coated surfaces appeared unaffected by the absence of CasL (**Fig 2B**). To quantitatively assess migration under these different conditions we employed an open source analysis software called Migrate3D ^54^. XY coordinates from individual cell tracks at each timepoint were imported into Migrate3D and analyzed for a variety of migration parameters. Initially, we evaluated the mean squared displacement (MSD) at every timepoint for control and CasL KO T cells migrating on either ICAM-1 and VCAM-1. As we and others have previously documented, T cell migration on ICAM-1 is significantly more robust than on VCAM-1 (**Fig 2 A-D** and ^21,25^). Importantly, we found that depletion of CasL had a significant effect on the MSD of T cells migrating on ICAM-1 coated surfaces, but not VCAM-1 coated surfaces (every timepoint **Fig 2C**, final timepoint **Fig 2E**). Loss of CasL negatively impacted several other important migratory parameters such as track length, speed, and straightness (**Fig 2F-H**). Again, these differences were only seen with T cells migrating on ICAM-1 coated surfaces but not VCAM-1 coated surfaces (**Fig 2F-H**). In addition to using our CRISPR/Cas9 system, we also repeated the above migration experiments using Cas^-/-^ T cells isolated from CasL germline knockout mice.

**Figure 2:**
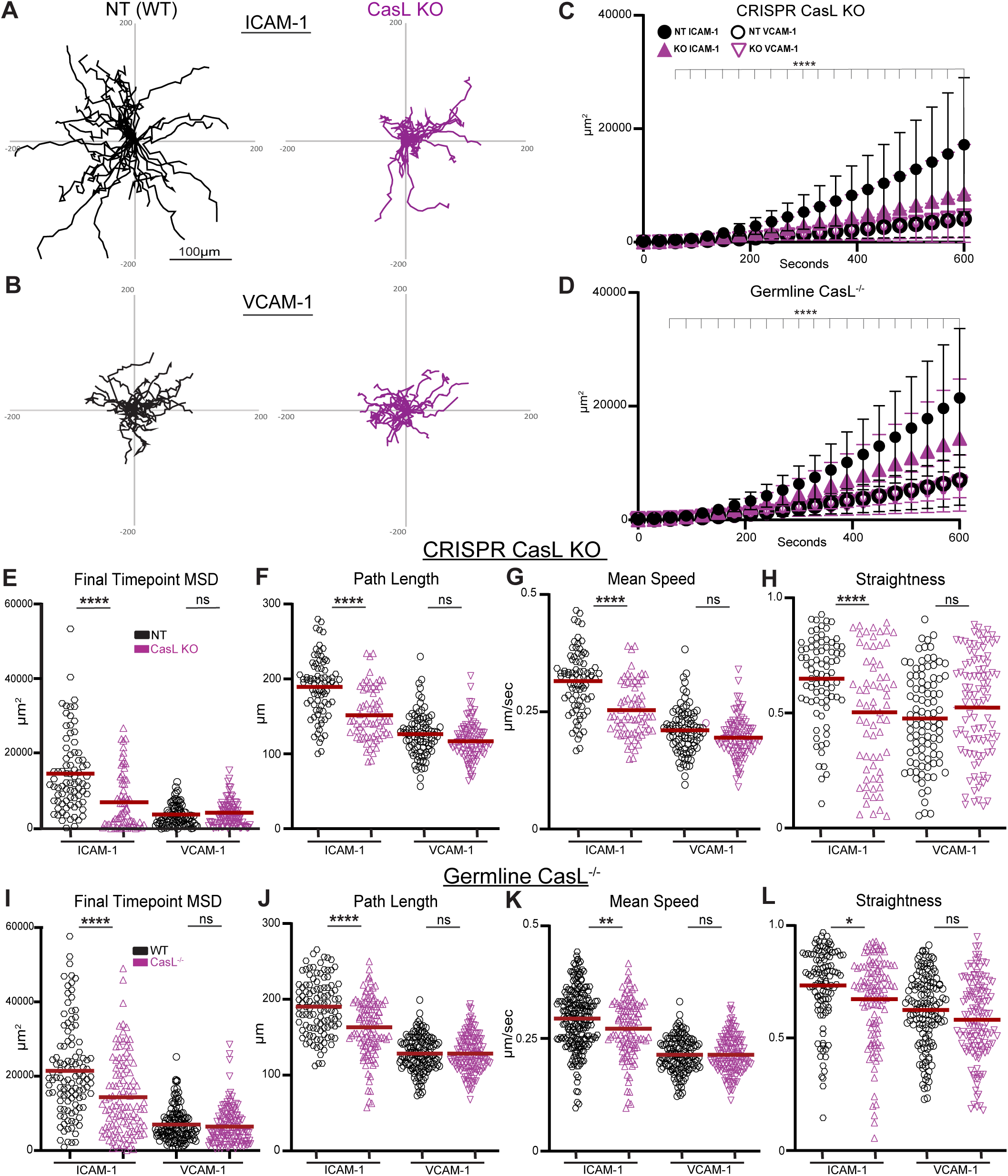
CasL is required for normal migration on ICAM-1 but not VCAM-1 coated surfaces. **(A)** Spider plots of individual cell tracks from 33 NT (WT) and 24 CasL KO T cells migrating on ICAM-1 and **(B)** from 25 NT and 25 CasL KO T cells migrating on VCAM-1, from a single representative experiment. **(C)** Mean squared displacement (MSD) over time from NT and CasL KO T cells migrating on ICAM-1 and VCAM-1 (n= 3 with 80 NT and 67 CasL KO T cells tracked on ICAM-1 and n=2 with 92 NT and 97 CasL KO T cells tracked on VCAM-1). **(D)** MSD over time from WT and germline CasL^-/-^ T cells migrating on ICAM-1 and VCAM-1 (n= 3 with 110 WT and 112 CasL^-/-^ T cells tracked on ICAM-1 and 137 WT and 135 CasL^-/-^ T cells tracked on VCAM-1). **(E)** MSD at final time-point, **(F)** path length, **(G)** mean velocity, and **(H)** straightness of NT and CasL KO T cells migrating on ICAM-1 and VCAM-1. **(I)** MSD at final time-point **(J)** path length **(K)** mean velocity and **(L)** straightness of WT and germline CasL^-/-^ T cells migrating on ICAM-1 and VCAM-1. T-test, * = p < 0.05, ** = p < 0.01, **** = p < 0.0001

CasL^-/-^ T cells also displayed migratory defects while migrating on ICAM-1 coated surfaces but not VCAM-1, consistent with the data using CRISPR/Cas9 to deplete CasL (**Fig 2D, and 2I-L**). Together, these data show that CasL specifically contributes to T cell migration on ICAM-1 but is dispensable for migration on VCAM-1.

### CasL restricts membrane blebbing in migrating T cells

During the low magnification imaging of cell migration on ICAM-1 we observed that CasL KO cells did not seem to spread to the surface and form clear lamellipodia at the same frequency as control cells. Indeed, when we quantified this observation we found that a greater proportion of WT cells spread to the surface compared to CasL KO T cells **(Supp Fig 2)**. To further characterize these apparent differences, we switched to 63x magnification and imaged WT and CasL^-/-^ T cells migrating on ICAM-1 coated surfaced for 1-2 minutes using high time resolution (1 frame-per-second). WT T cells migrating on ICAM-1 formed prominent lamellipodia and moved smoothly with continuous sheet-like projections, as previously described ^15,17,21,23,24,55^ (**Fig 3A and Supp movie 1**). In contrast, CasL^-/-^ T cells rarely formed a stable lamellipodium but instead formed numerous aberrant membrane protrusions in the front and along the sides of the cell (**Fig 3A and Supp movie 2**). In many cases, smooth projections appeared to pop from the membrane rapidly, usually within 2 seconds or less, indicative of membrane blebs (as reviewed in ^56^). Imaging CasL^-/-^ T cells expressing lifeact-GFP confirmed that these structures were initially devoid of F-actin, consistent with the architecture of membrane blebs (**FIG 3B and Supp movie 3**). To quantify the extent of blebbing in migrating T cells, we counted blebs on a per cell basis for the duration of the imaging period. Protrusions were only classified as blebs if they appeared in under 2 seconds, and displayed a smooth, rounded appearance. Remarkably, roughly 50% of cells lacking CasL displayed at least one membrane bleb during a one-minute period, while less than 10% of control cells blebbed during the same timeframe (**Fig 3C-D**). In addition, control T cells that blebbed generally only displayed one bleb in a one-minute period, while T cells lacking CasL displayed numerous blebs, sometimes over 10 in one minute (**Fig 3C-D**). We obtained similar results using both genetic systems to ablate CasL levels, CRISPR/Cas9 and germline CasL^-/-^ (**Fig 3C-D**). Taken together, these results indicate that CasL promotes normal leading-edge morphology and restricts membrane blebbing in T cells migrating on ICAM-1.

**Figure 3:**
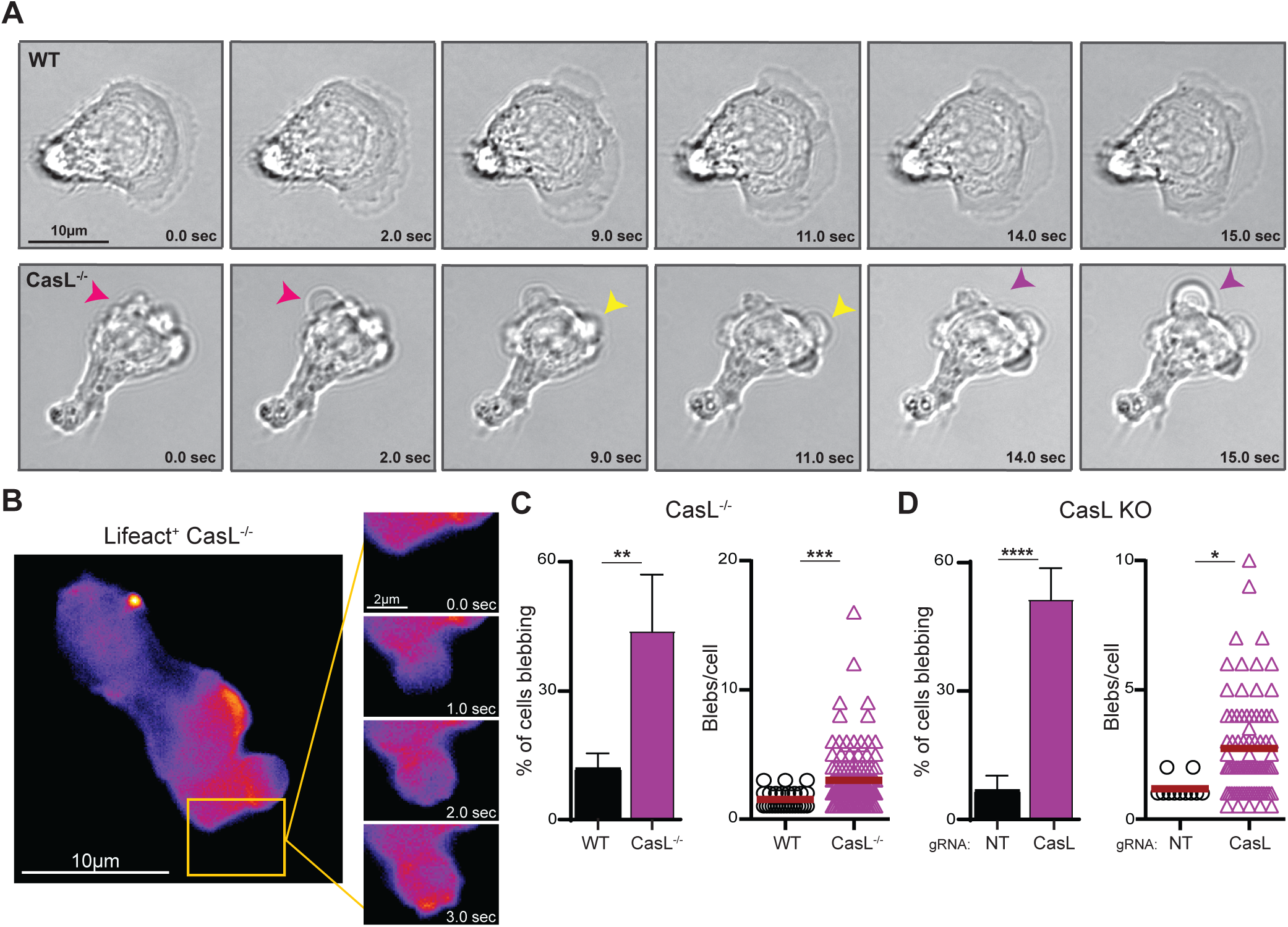
CasL restricts membrane blebbing in migrating T cells. WT and germline CasL^-/-^ CD4^+^ T cells were allowed to migrate on ICAM-1 coated surfaces and imaged at 63x for 1-2mins with 1 second between frames. **(A)** individual frames from WT (upper panel) and CasL^-/-^ (lower panel). Darts indicate the rapid appearance of blebs in the CasL^-/-^ T cells. **(B)** CasL^-/-^ T cells expressing lifeact-GFP were imaged at 63x for 1min with 1 second timepoints. **(C)** Quantification of blebbing in WT and CasL^-/-^ T cells migrating on ICAM-1 (n=3, 229 WT and 262 CasL^-/-^ cells). **(D)** Quantification of blebbing in NT or CasL KO T cells migrating on ICAM-1 (n=3, 150 NT and 148 KO cells). T-test, * = p < 0.05, ** = p < 0.01, *** = p < 0.001, **** = p < 0.0001

### Y-27632 treatment inhibits blebbing in T cells lacking CasL

Major causes of blebbing include dysregulated actomyosin contractility and/or altered cortex integrity ^21–27^. The Rho-associated kinase (ROCK) inhibitor Y-27632 (Y-27) is commonly used to globally perturb actomyosin contractility and can inhibit membrane blebbing in a variety of contexts, likely via decreasing cell contraction and relieving cytosolic pressure (for example ^57–59^). To test if the increased blebbing in CasL KO T cells is sensitive to ROCK inhibition, we repeated the above imaging experiments in the presence of the Y-27. When T cells lacking CasL were treated with Y-27 (2.5µM), the blebbing immediately ceased and, astonishingly, they formed lamellipodia indistinguishable from control T cells (**Fig 4A-C, Supp movie 4 and 5**). These results suggest lack of CasL may perturb RhoA/ROCK signaling or one of the downstream effectors of this pathway.

**Figure 4:**
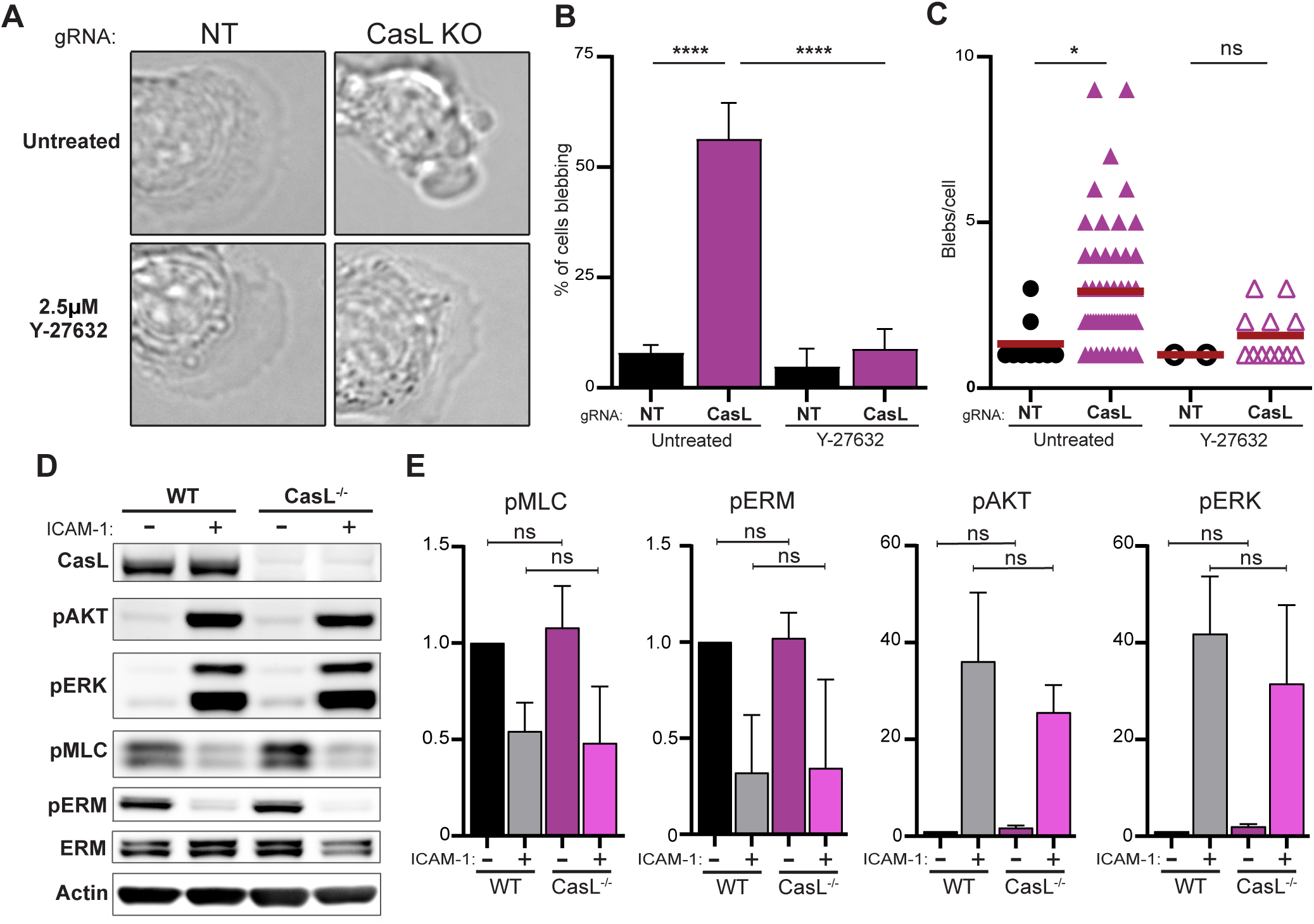
Y-27632 treatment abolishes blebbing in T cells lacking CasL. **(A)** Representative images of the front of migrating NT and CasL KO T cells before and after treatment with 2.5µm Y27632. **(B and C)** Quantification of blebbing in NT and CasL KO T cells before and after treatment with 2.5µm Y27 (n=3 with 103 NT T cells before and 56 cells after the addition of Y27, 108 CasL transduced T cells before and 67 cells after the addition of Y27). One-way ANOVA test, * = p < 0.05, **** = p < 0.0001 **(D)** WT and CasL^-/-^ CD4^+^ T cells were allowed to adhere to ICAM-1 coated surfaces for 20 mins, lysed, and immunoblotted with the indicated antibodies. **(E)** Quantification of pAKT, pERK, pERM, and pMLC band intensity from n=4. T-test, * = p < 0.05, **** = p < 0.0001

Therefore, we decided to biochemically probe the activation state of important cytoskeleton regulators that are associated with ROCK signaling and implicated in blebbing, myosin light chain and ezrin-radixin-moesin (ERM) proteins, in T cells lacking CasL. To do this, we serum starved control and CasL^-/-^ T cells then allowed them to migrate on ICAM-1 coated surfaces, or incubated them in suspension as an unstimulated control. After 20 minutes, T cells were lysed and immunoblotted with the indicated phospho-specific antibodies. ICAM-1 stimulation caused a global decrease of both pMLC and pERM in WT T cells, suggesting the overall relaxing of contractility and membrane tension and upon cell spreading on ICAM-1 (**Fig 4D-E**). Surprisingly, despite the abnormal blebbing in CasL^-/-^ T cells, there were no significant differences observed in the levels of phosphorylated myosin light chain or phosphorylated ERM proteins with or without ICAM-1 stimulation compared to WT T cells, indicating CasL does not influence the global activity of these cytoskeletal regulators (**Fig 4D-E**). In addition, we probed two major signaling pathways induced by ICAM-1 that can affect T cell migration, PI3K and MAPK, by immunoblotting for pAKT and pERK, respectively. T cells lacking CasL robustly activated these pathways similar to WT T cells (**Fig 4D-E**). These data indicate that the abnormal blebbing observed in T cells lacking CasL is likely not a result of global changes in pMLC or pERM, and that these T cells can still form productive interactions with ICAM-1 surfaces and successfully trigger important signaling events. However, Y-27 treatment abolished blebbing in CasL^-/-^ T cells, suggesting at least some involvement of hyper contractility or cytoskeletal dysregulation in the aberrant blebbing seen in CasL^-/-^ T cells.

### CasL controls the balance of cytoskeletal components in migrating T cells

Our observations from live cell imaging showing that CasL deficient T cells lacked a well-spread leading edge but instead formed numerous membrane blebs and irregular projections (**Fig 3-4**) suggested that CasL may control the normal cytoskeletal architecture in migrating T cells. To determine if CasL influences the spatial distribution of major cytoskeletal components, WT and CasL^-/-^ T cells were allowed to migrate on ICAM-1 coated surfaces, fixed, and then stained with phalloidin to label F-actin, along with an antibody that recognizes the active, phosphorylated version of myosin light chain (S19). WT T cells displayed the canonical actin rich leading edge that is generally observed in T cells migrating on ICAM-1 (**Fig 5A**). However, CasL^-/-^ T cells exhibited far less F-actin at the cell front (**Fig 5A**). In addition, compared to WT T cells, CasL^-/-^ T cells showed an apparent accumulation of pMLC in the rear of the cell, in many cases directly behind the nucleus (**Fig 5A**). To quantify these observations, we first outlined each cell and made cell area and fluorescence intensity measurements for each channel on a cell-by-cell basis. We excluded round cells to ensure we were only focusing our analysis on migratory cells. Using these criteria, we first found that CasL^-/-^ T cells had a significantly smaller cell area than their WT counterparts (**Fig 5C**), suggesting CasL affects cell spreading to the surface. In addition, as visually observed in **Fig 5A**, CasL^-/-^ T cells also displayed significant decreases in both total and mean F-actin signal per cell (**Fig 5D-E**). Total pMLC per cell was also significantly decreased, however, there were only very modest decreases in mean intensity of pMLC (**Fig 5F-G)**. These data suggest that difference in total pMLC in CasL^-/-^ T cells likely stems from the decrease in cell area of these T cells **(Fig 5C)**, not intrinsic differences in total amount of pMLC per cell, consistent with the immunoblot experiments in **Fig 4D-E**. However, there did appear to be a specific accumulation of pMLC in the rear of CasL^-/-^ T cells (**Fig 5A and Supp Fig 3**), therefore, to quantify the relative distribution of pMLC in migrating T cells, we segmented each cell into a front and back and calculated the percent of signal from each compartment (**Fig 5B**). Using this analysis we found that, indeed, there was a significant accumulation of pMLC in the back half of CasL^-/-^ T cells compared to WT T cells (**Fig 5H**). This abnormal accumulation correlated with what appeared to be sites of nuclear deformation indicating that the nucleus may be under pressure and getting squeezed forward (**Fig 5A and Supp Fig 3**). Indeed, when we quantified nuclear circularity, nuclei from CasL^-/-^ T cells were significantly more deformed (**Fig 5I**). Together, these data show that lack of CasL leads to a decrease in F-actin in the front of migrating T cells accompanied by an abnormal accumulation of pMLC behind the nucleus.

**Figure 5:**
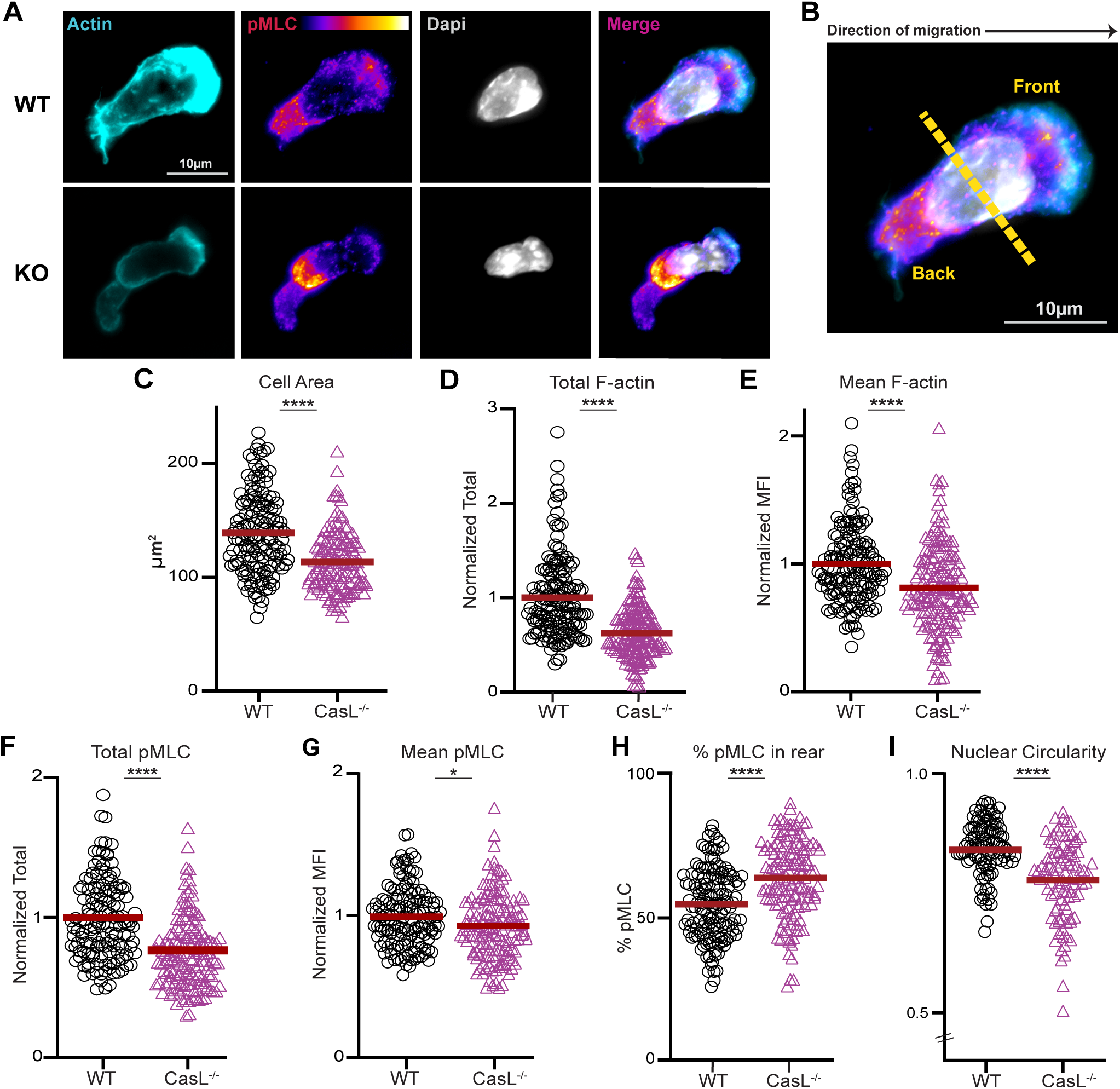
CasL controls the balance of cytoskeletal components in migrating T cells. **(A)** WT and CasL^-/-^ CD4^+^ T cells were allowed to migrate on ICAM-1 for 20mins, fixed, and stained with anti-pMLC, florescent phalloidin, and DAPI. **(B)** Example of demarcation line showing how measurements from the front and back of individual cells were quantified. **(C)** cell area, **(D)** total actin per cell, **(E)** mean actin per cell, **(F)** total pMLC per cell, **(G)** mean pMLC per cell, **(H)** percent of pMLC in cell rear, and **(I)** nuclear circularity. N=3, 143 WT and 149 CasL^-/-^ T cells. T-test, * = p < 0.05, **** = p < 0.0001

### CasL promotes migration into inflamed tissue *in vivo*

Our *in vitro* work strongly suggest that CasL is an important factor for integrin mediated migration, however it is unclear how CasL contributes to T cell migration in a more complex inflammatory *in vivo* setting. Therefore, we utilized a competitive *in vivo* trafficking assay to determine if CasL is needed for T cell migration to peripheral tissues and/or lymphoid organs during inflammation. The assay is based on a mouse model of graft-versus-host disease (GvHD) in which T cells are adoptively transferred into an MHC mismatched recipient and proceed to migrate into and attack peripheral tissues **(Fig 6A)**. Briefly, recipient BALB/c (H2K^d^) mice were irradiated and received T cell-depleted bone marrow from B6 mice (H2K^b^), along with a 50/50 mixture of WT/CasL^-/-^ T cells, or WT/WT T cells as a control. Congenically labeled WT donor T cells (CD45.1^+^) were used in each group to differentiate between the competing CasL^-/-^ T cells (CD45.2^+^) or the competing WT T cells (CD45.2^+^) in the control group. Before initial injections, flow cytometry was run on both test group bone marrow and T cell mixes to confirm near 50/50 input ratios for CD45.1^+^ to CD45.2^+^ cells **(Fig 6B)**. After 7 days, mice were euthanized and the spleen, lymph nodes, liver, and lung tissues were collected and stained for flow cytometry. T cells were gated to include only donor T cells, then the percentages of CD45.1^+^ (WT) to CD45.2^+^ (WT or CasL^-/-^) were determined **(Fig 6C)**. Using this strategy, we observed no difference in migration of CasL^-/-^ T cells into either the spleen or lymph nodes compared to WT T cells **(Fig 6D,E)**. However, CasL^-/-^ T cells were outcompeted by their WT counterparts in the liver and lungs, showing that T cells lacking CasL have impaired migration to these locations **(Fig 6F,G)**. Collectively, these data indicate CasL is an important factor in promoting T cell migration into inflamed tissue but dispensable for trafficking into lymphoid organs.

**Figure 6:**
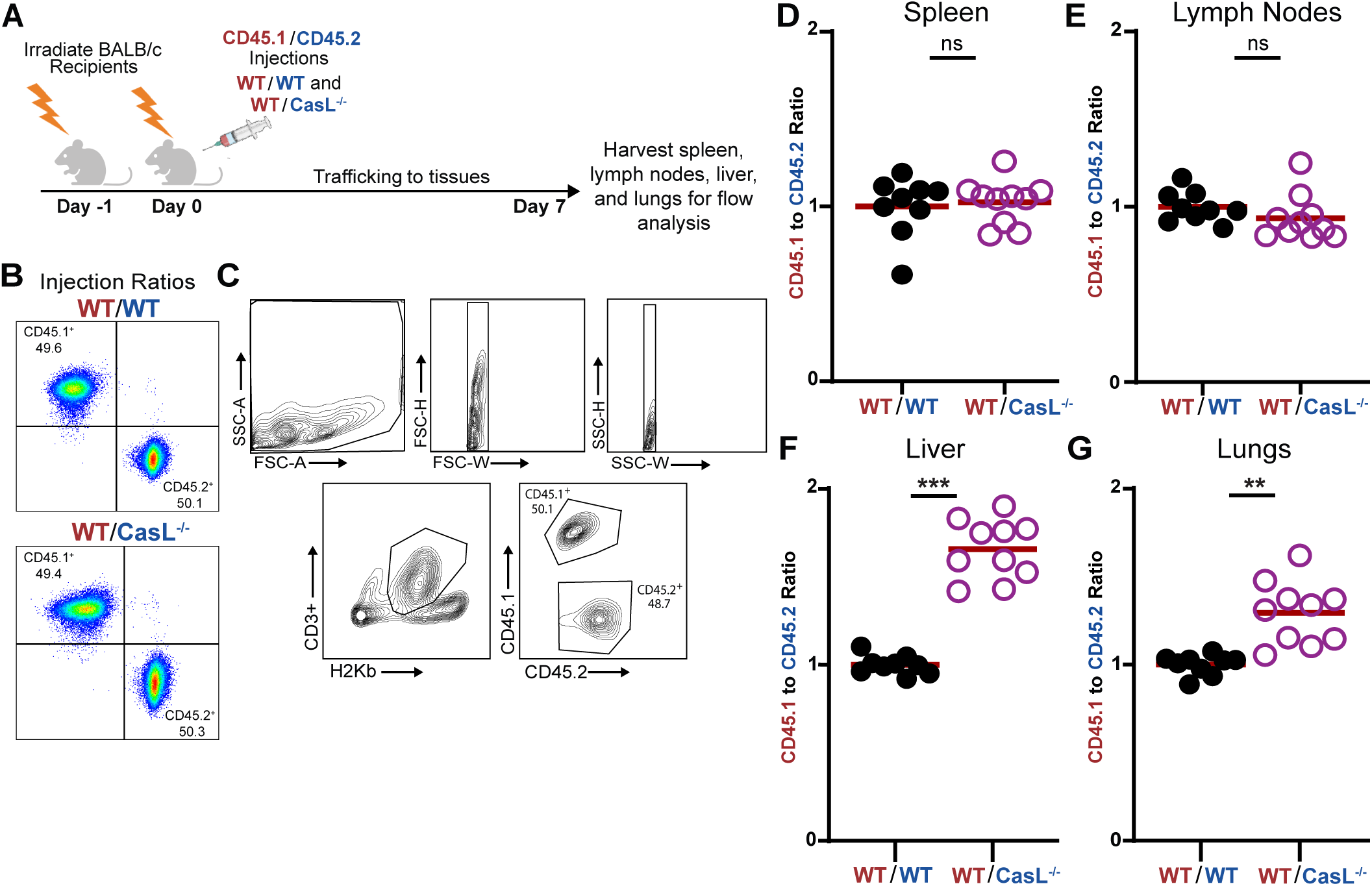
CasL promotes migration into inflamed tissue. **(A)** Murine model of GvHD used to compare migration of adoptively transferred WT and CasL^-/-^ T cells into different tissues. **(B)** Flow cytometry of input ratios of CD45.1^+^ (WT) and CD45.2^+^ (WT or CasL^-/-^) T cells prior to transplant. **(C)** Gating strategy to identify CD45.1^+^ and CD45.2^+^ T cell populations from spleen, lymph node, liver, and lung tissues. Ratio of CD45.1^+^ to CD45.2^+^ T cells collected from the **(D)** spleen, **(E)** lymph nodes, **(F)** liver, and **(G)** lungs of recipient Balb/C mice (n= 9 WT/WT mice and 10 WT/ CasL^-/-^ mice). T-test, ** = p < 0.01, *** = p < 0.001

## Discussion

Using two genetic systems to ablate CasL in primary T cells, we found that CasL was specifically required for efficient migration on ICAM-1 (LFA-1 dependent migration), but not on VCAM-1 (VLA-4 dependent migration) coated surfaces. Surprisingly, CasL was needed to maintain normal leading edge integrity during migration, as T cells lacking CasL displayed numerous, uncontrolled membrane blebs and migrated more erratically. *In vivo*, T cells lacking CasL could enter and expand normally in secondary lymphoid organs, but had diminished migration into inflamed peripheral tissue. Together, these data indicate CasL is a major scaffold that links LFA-1 engagement to cytoskeletal responses in migrating T cells.

Our finding that CasL influences migration on ICAM-1 but not VCAM-1 coated surfaces reinforces the idea that these two integrins elicit very different migratory responses. T cells migrating on ICAM-1 display a more broad, stable, actin rich front, while T cells migrating on VCAM-1 use more discrete pseudopodia and migrate slower with less directionality ^21^. We and others have previously shown that both ICAM-1 and VCAM-1 support strong adhesion ^21,60^, therefore the differences observed in migration are not due to lower adhesion of T cells to VCAM-1, but likely a difference in downstream signaling initiated by B2 or B1 integrins, in this case LFA-1 and VLA-4, respectively. While T cells do not form canonical focal adhesions, it was recently demonstrated that they do indeed form small, dynamic focal adhesion-like complexes that contain talin and vinculin in response to both B2 and B1 ligands ^61^. Because CasL is known to localize to focal adhesions in other cell types ^34–39^, it is possible CasL is an actual component of specific adhesion complexes initiated by B2, but not B1 integrins. Interestingly, the presence of CasL did not seem to affect LFA-1 signaling *per se*, as other important pathways like PI3K (which is known to regulate cell migration) remained unchanged in CasL KO T cells (**Fig 4**). These data suggest CasL signaling proceeds parallel to other major pathways or represents a branch point in signaling that controls only cytoskeletal responses.

Close inspection of CasL KO T cells migrating on ICAM-1 revealed that instead of forming a controlled actin-rich cell front like WT T cells, T cells lacking CasL formed numerous membrane blebs. While T cells have been hypothesized to use bleb-based migration in certain situations ^62^, blebbing is rare during integrin mediated migration (**Fig 3-4**). Importantly, the blebs observed in T cells lacking CasL were small, transient blebs, and not the larger stable blebs seen in some cell types ^56^. These small blebs generally form due to disruption of membrane-cortex integrity to the point where cytosolic pressure can force detachment of the membrane from the underlying cortex ^56,63,64^. This mechanism is consistent with our data showing that T cells lacking CasL have a clear decrease of F-actin in the front of migrating T cells (**Fig 5**), although we did not visualize the cortex directly. Knocking down Arp3 in T cells (to disrupt the Arp2/3 complex) also resulted in uncontrollable blebbing ^65^, supporting the idea that changes in membrane proximal actin levels or organization can lead to a biophysical environment conducive to blebbing. In addition, CasL KO T cells also showed a clear increase of pMLC in the rear of migrating cells, indicating contractile forces are being generated. Importantly, the accumulation of pMLC looked to be deforming the nucleus, pushing it toward the cell front, a process known to increase cytosolic pressure ^66^. Relaxing contractile forces with Y-27 treatment reversed the aberrant blebbing observed in CasL KO T cells, suggesting the nuclear squeezing coupled with decreases in anterior F-actin likely account for blebbing in T cells lacking CasL.

There are several potential mechanisms of how CasL could function to help maintain normal cytoskeletal architecture and cell shape during T cell migration. Given the perturbations in both actin and myosin in migrating CasL KO T cells, it is likely CasL helps to maintain the normal activity of one or more of the Rho family GTPases that define the cytoskeletal activity at the front (Rac and Cdc42) or the rear (RhoA) ^67–69^.

Indeed, a study in B cells showed CasL interacts with AND-34, a Cdc42 GEF that controls actin structure ^70^. CasL could physically localize near or within adhesion sites, as mentioned above, and help recruit specific GEFs and/or GAPs to influence GTPase activity. The 13 putative pTyr sites on CasL provide considerable scaffold space for complex signaling regimes to arise. Interestingly, a recent pre-print revealed a mechanical feedback mechanism between GTPases that helps establish front/back polarity in immune cells ^71^. Our data suggests CasL may affect membrane/cortex integrity, therefore it is possible CasL is involved in maintaining the cortical actin structure needed for proper mechanical feedback, either by acting as a signaling scaffold or as a structural component of the cortex itself. Unfortunately, we have not found a suitable antibody to reliably determine the localization CasL in migrating T cells. Future work will aim to answer some of the outstanding questions about how CasL regulates cytoskeletal architecture in T cells.

Previous work in CasL^-/-^ germline knockout mice showed that at steady-state there were fewer lymphocytes in secondary lymphoid organs compared to WT mice ^50^. However, when CasL^-/-^ T cells were adoptive transferred into WT mice, there was only a very modest decrease of T cell migration into the spleen and lymph nodes, indicating the larger differences observed at steady-state may be due to compounding factors from the germline knockout mice used ^50^. T cell migration under inflamed conditions, were integrin integrins are particularly important, was not evaluated. Using an allogeneic hematopoietic transplant model that mimics graft-versus-host disease, we now show that CasL is specifically needed for migration into inflamed peripheral tissue. CasL^-/-^ T cells expand and inhabit secondary lymphoid organs normally but were outcompeted by WT T cells roughly 1.75 to 1 in the liver, and 1.25 to 1 in the lung. The selective defect in migration to inflamed tissue may represent a specific requirement for integrin-dependent cytoskeletal changes to help squeeze through the vascular barrier versus the relatively easy passage of lymphocytes into inflamed secondary lymphoid organs. These findings highlight the importance of integrin signaling for T cell migration into inflamed tissue and open the door for future work is aimed at determining which signaling and cytoskeletal events downstream of CasL specifically control migration into different inflamed tissues.

## Materials and Methods

### Antibodies and reagents

Anti-MHCII clone M5/114, anti-CD8 clone 2.43, anti-CD3 clone 2C-11, and anti-CD28 clone PV-1 were obtained from BioXCell. CD90.2 magnetic beads used to isolate bone marrow and T cells with LD and LS columns were purchased from Miltenyi Biotec. Western blot and immunofluorescence antibodies anti-pAKT(S743) clone D9E (cat#4060S), anti-Ezrin/Radixin/Moesin (cat#3142S), anti-pEzrin(Thr567)/Radixin(Thr564)/Moesin(Thr557) (cat#3141S), anti-pERK/P-p44/42 MAPK (cat#9101S), anti-actin clone 13E5 (cat#4970S), and anti-pMyosin light chain (S19) (cat#3671S) were all purchased from Cell Signaling. Anti-CasL antibody clone 2G9 was produced in house via hybridoma and purified using protein-G. Secondary antibodies conjugated with appropriate fluorophores were obtained from Jackson Immunoresearch or Life Technologies. Pre-conjugated flow cytometry antibodies anti-CD3/PerCP_Cy5.5 (cat#100217), anti- CD4/PE (cat#100407) or PE/Cy7 (cat#100421), anti-CD8/FITC (cat#126605), anti-ß1/APC (cat#102215), anti- ß2/PE (cat#101407), anti-CD45.1/APC (cat#110713), and anti-CD45.2/APC_Cy7 (cat#109823) were all purchased from BioLegend. All antibodies used in this study were validated by the manufacturer. Dapi was obtained from Sigma Aldrich. Recombinant mouse ICAM-1 and VCAM-1 were purchased from R&D Systems. The ROCK inhibitor Y-27632 was purchased from Cayman Chemicals.

### Mice

All mouse husbandry was done in the Upstate Medical University mouse facility in accordance with the Institutional Animal Care and Use Committee guidelines. The CasL germline knockout mice (C57BL/6J background) were originally created by Sachiko Seo and Hisamaru Hirai ^50^, and kindly provided to our lab by Erica Golemis (Fox Chase Cancer Center). WT B6 (C57BL/6J, H-2K^b^, strain #000664), Cas9 (B6J.129(Cg)- *Igs2^tm1.1(CAG-cas9*)Mmw^*/J, strain #028239), Thy1.1^+^ (B6.PL-*Thy1^a^*/CyJ, strain #000406) and CD45.1^+^ (B6.SJL- *Ptprc^a^ Pepc^b^*/BoyJ, H-2K^b^, strain #002014) mice were purchased from The Jackson Laboratory. BALB/C (H- 2K^d^, strain #027) mice used as recipients in the competitive trafficking assay were purchased from Charles River. All mice were 8-12 weeks of age when cells were harvested and both male and female mice were used equally as sources of T cells for *in vitro* assays.

### Cell culture

For *in vitro* assays, primary mouse CD4^+^ T cells were isolated from spleens and lymph nodes from 8–12 week old mice. Briefly, tissues were homogenized and filtered before incubation with ACK lysis buffer (77mM NH_4_Cl, 5mM KHCO_3_, and 55µM EDTA pH 7.2-7.4) for 1 min to remove red blood cells (RBC). After RBC lysis, cells were washed and incubated with anti-CD8^+^ and anti-MHCII for 20min at 4°C. After initial incubation, cells were washed and mixed with BioMag Goat anti-rat IgG beads (Qiagen cat#310107) and incubated an additional 15min at 4°C before samples were placed onto a magnetic rack for three rounds of magnetic separation. Isolated CD4^+^ T cells were then counted and immediately activated on a 24-well plate coated with 5µm/mL anti-CD3 and 2µm/mL anti-CD28 at 1 X 10^6^ cells per well. Activation was done in complete T cell medium composed of Corning DMEM supplemented with 5% FBS, 1% each penicillin/streptomycin, non-essential amino acids, and Glutamax (all purchased from Gibco), plus 2µL 2-mercaptoethanol (obtained from Sigma Aldrich). After 48hrs of activation, T cells were removed from activation and mixed at a 1:1 volume ratio with complete T cell medium containing recombinant IL-2 (obtained from Sigma Aldrich) to a final concentration of 20U/mL IL-2. T cells were used on days 5-6 post-harvest for all experiments. 293T cells (for retroviral production) were maintained in DMEM supplemented with 10% FBS, 1% non-essential amino acids (Gibco), and Glutamax (Gibco) and regularly screened for mycoplasma.

### Retroviral production

Specific gRNAs to CasL were cloned into the retroviral transfer vector described in Huang *et al*. 2019, MRIG ^51^. This vector was previously modified to express a puromycin resistance gene in place of the original GFP for downstream selection ^21^. The gRNAs sequences used in this study were as follows: non-targeting (NT) 5’-GCGAGGTATTCGGCTCCGCG-3’; CasL (sequence 1) 5’-AGTAGGCTGCTCATGACCGG-3’; CasL (sequence 2) 5’-GGAATGTCATATACCCCTTG-3’. To produce retroviral particles containing guide RNAs to CasL, 293T cells (originally obtained from ATCC) were grown to 70% confluency before being co-transfected with 4.5µg MRIG-puro along with 3.5µg pCL- eco packaging vector (Addgene #12371). Transfection was done using calcium phosphate purchased from Invitrogen (cat# K278001). Transfection mixes remained on 293T cells overnight before being replaced with fresh media, after which cells were incubated for an additional 24- 30hrs. Retroviral containing supernatants were collected and centrifuged for 10min at 1000*g* to remove cellular debris, and polybrene was added to clarified viral supernatants at a final concentration of 8µg/mL. Retroviral supernatants were then immediately applied to T cells for transductions.

To produce lifeact-GFP retrovirus, the transfer vector pMSCV2.1 containing the sequence for lifeact-GFP (a kind gift from Janis Burkhardt) was co-transfected with pCL- eco packaging vector exactly as above.

### T cell transduction

CD4^+^ T cells isolated from Cas9-expressing mice were activated on 24-well plates coated with 5µg/mL of anti-CD3 and 2 µg/mL anti-CD28 at 1x10^6^ cells per well in T cell complete medium. After 24hrs of activation the T cell media was collected and reserved for use post-transduction, and 800uL of viral supernatants were gently added per well before plates were incubated at 37°C for 10min. Following incubation, plates were centrifuged at 1100*g* for 2hrs at 35°C. Directly following spinoculation, plates were incubated at 37°C for 10min and then viral supernatants were removed and replaced with 1.5mL of conditioned media mixed with fresh complete T cell medium at a 2:1 volume ratio. T cells were cultured for an additional 24hrs before being removed from activation and mixed at a 1:1 volume ratio with complete T cell medium containing recombinant 40 U/mL IL-2 to create a final concentration of 20U/mL. After 2hrs incubation in IL-2 media at 37°C, media containing puromycin was added so that the final concentration was 4µg/mL. Transduced cells were used for migration assays after 3 days of selection in puromycin (day 5 post-isolation).

To generate lifeact-GFP T cells expressing CasL, CD4^+^ T cells from CasL^-/-^ mice were activated and transduced exactly as above, with the expectation that no puromycin was added during T cell expansion.

### Transwell assay

CD4^+^ T cells were serum starved in DMEM without FBS for 2hrs at 37°C. After serum starve, 5.0 x 10^5^ – 1.0 x 10^6^ cells were allowed to settle on transwell inserts with 3µm pores (Corning, ref# 3415) for 15min before filling the bottom chamber with either FBS-free media as a control or media supplemented with 1% FBS as a migratory stimulus. After a 2hrs migration period at 37°C, transwell inserts were removed with one gentle wash on the bottom before cells from each well were manually counted using a hemocytometer. The percentage of cells that successfully transmigrated through 3µm inserts was calculated based on the number of cells initially plated.

### Biochemical analysis of T cell signaling in response to surface bound ICAM-1

60-mm tissue culture dishes (Corning, 430166) were coated with 2µg/mL ICAM-1 overnight at 4°C. Activated CD4^+^ T cells were serum starved in FBS free DMEM for 2hrs at 37°C before being washed and resuspended in warm L15 medium (Gibco) supplemented with 2mg/mL D-glucose. Cells were incubated at 37°C for 10min before 10 x 10^6^ cells were added to 60mm dishes and allowed to interact with surface bound ICAM-1 for 20min, or left unstimulated as a control. ICAM-1 stimulated cell lysates were collected by placing dishes on ice, aspirating off L15 media, and adding 1.25x cold lysis buffer (1x composition: 1% Triton X-100, 150mM NaCl, 50mM Tris-HCl pH 7.5, 5mM NaF, 1mM sodium orthovanadate, and Roche EDTA-free protease inhibitor). Unstimulated control cells were lysed in suspension by placing on ice and adding cold 2x lysis buffer at 1:1 volume ratio. Lysates were kept on ice and vortexed periodically for 15min, followed by centrifugation at 15,000*g* for 10min at 4°C. Clarified whole cell lysate was mixed with 4x sample buffer (Invitrogen) containing DTT (Roche, 50mM final concentration) and heated for 8min at 95°C before separation with SDS-PAGE.

### Western blotting

Proteins were separated by SDS-PAGE using the Invitrogen Novex Mini-cell system with NuPAGE 4-12% BisTris gradient gels. Proteins were transferred to nitrocellulose membranes (0.45µm, BioRad) and blocked using Li-Cor blocking buffer mixed at a 1:1 volume ratio with PBS for 1hr at room temperature. Primary antibodies were mixed in TBST (50mM Tris, 150mM NaCl, 0.1% tween-20) with 2% BSA and added to membranes for overnight incubation at 4°C. Primary antibody dilutions were as follows: anti-CasL (1:100), anti-pAKT (1:500), anti-pERK (1:500), anti-pMLC (1:500), anti-pERM (1:500), anti-ERM (1:1000), and anti-actin (1:5000). Membranes were washed three times with TBST for 10 minutes each wash before the addition of fluorophore-conjugated secondary antibodies diluted in TBST with 2% BSA and incubated for 1hr at room temperature. Membranes were then washed three times with TBST and imaged using a ChemiDoc Imaging System (Biorad). Band intensity measures were done using ImageJ software.

### Imaging T cell migration on ICAM-1 and VCAM-1

Ibidi 8-well, glass-bottomed chamber slides were coated with ICAM-1 or VCAM-1 at 2µg/mL overnight at 4°C. Activated CD4^+^ T cells were washed and resuspended in Leibovitz’s L-15 medium (Gibco) containing 2mg/mL D-glucose before 5 x 10^4^ cells were added to each well. Slides were incubated at 37°C for 20min before being gently washed once to remove non-adherent cells and debris, then mounted for imaging. Imaging was performed on a Marianas system (3i) consisting of a Ziess Axio Observer 7 equipped with a X-Cite mini+ light source (Excelitas) and a Prime BSI Express CMOS camera (Photometrics) enclosed in an environmental chamber (Okolab). Time-lapse images were collected every 30 seconds for 10min using a 10x or 20x objective at 37°C with SlideBook software (3i) and exported to ImageJ for analysis. Individual cells were tracked using the manual tracking plugin to generate XY coordinates for all motile cells. Cells were considered motile if they moved more than 2 cell-lengths during the entire imaging period. We also only included cells that stayed in the field-of-view for the entire imaging period. XY coordinates were analyzed using the open-source software Migrate3D ^54^ to determine a variety of migratory parameters, including mean squared displacement, track length, Euclidean distance/displacement, speed, and directionality.

To quantify blebbing, T cells migrating on ICAM-1 as described above were imaged using a 63x objective with 1 second time intervals for 1-2mins total. Individual cells were monitored frame-by-frame for the appearance of membrane blebs which were classified as smooth, rounded protrusions that appeared to “pop” the cell membrane out in 2 seconds or less.

### Quantification of pMLC and F-actin in migrating T cells

Ibidi 8-well, glass-bottomed chamber slides were coated with ICAM-1 at 2µg/mL overnight at 4°C. Activated CD4^+^ T cells were washed and resuspended in L-15 medium containing 2mg/mL D-glucose and 5 x 10^4^ cells were added per well. Slides were then incubated at 37°C for 25min, followed by fixation with 3.7% paraformaldehyde (Electron Microscopy Sciences). Cells were blocked and permeabilized (1% BSA and 0.15% Triton X-100 in PBS) for 15min at room temperature then incubated with anti-phospho myosin S19 (in perm/block solution) overnight at 4°C. Slides were washed three times with perm/block solution and incubated with secondary antibodies, DAPI, and fluorescent phalloidin, at room temperature for 45min. Slides were then washed three times with perm/block solution and put in PBS for imaging. To quantify F-actin and phosphorylated myosin light chain, 0.5µm *z*-stacks were collected from the bottom of the cell moving upward through the cell a total of 8µm. Each z-slice was background subtracted and the image was projected.

Individual cells were manually outlined, and total intensity of each fluorescence channel was measured. In addition, cells were divided in half using the nucleus as a guide to calculate front-back ratios for fluorescence channels. All analysis and intensity measures were done using SlideBook 2023.

### In vivo competitive migration

Bone marrow was collected from the femurs and tibia of WT B6 H-2K^b^, Thy1.1^+^ mice (B6.PL-*Thy1^a^*/CyJ, strain #000406) and depleted of T cells using anti-CD90.2 magnetic beads (Miltenyi). T cells were collected from the spleens of WT C57BL/6J mice (CD45.2^+^), CasL^-/-^ mice (CD45.2^+^), and WT CD45.1^+^ congenically labeled mice (B6.SJL-*Ptprc^a^ Pepc^b^*/BoyJ), using a CD90.2 positive selection. Recipient BALB/C mice (H-2K^d^) were irradiated in two sessions for 170 seconds each at 160kV, with the initial irradiation roughly 18hrs before transplant injections. Within 4hrs of the second irradiation, recipients were injected with either bone marrow alone, or bone marrow supplemented with a 50/50 ratio of WT/WT(CD45.1^+^) or CasL^-/-^ /WT (CD45.1^+^). Before injections, the input ratio of CD45.2^+^/CD45.1^+^ T cells in both test groups injection mixtures was confirmed by flow cytometry. Test mice were given with roughly 2-3 million T cells (1-1.5 million each CD45.2^+^ and CD45.1^+^) mixed into bone marrow injections. 7 days after bone marrow transplants, mice were euthanized and spleens, lymph nodes, livers, and lungs were harvested and processed for flow cytometery.

### Flow cytometry

For *in vivo* migration experiments, cells were initially stained with Live/Dead Zombie Aqua (BioLegend cat #423101) in PBS following the manufacturer’s protocol. Antibodies were diluted at 1:200 in ice cold FACS buffer (PBS, 1% BSA, and 0.02% EDTA), and cells were stained for 1h followed by 2 washed in ice cold FACS buffer. Antibodies used were; anti-H2Kb, anti-CD3, anti-CD4, anti-CD8, anti-CD45.1, and anti-CD45.2. Flow cytometry was performed using a Fortessa (BD Biosciences) cytometer and data was analyzed using FlowJo software (FlowJo LLC).

To confirm expression of both LFA-1 and VLA-4 on CasL deficient T cells, cells were stained with Samples were stained with, anti-CD3, anti-CD4, anti-CD8, anti-ß1, and anti-ß2.

### Statistical analysis

Statistics were calculated using GraphPad Prism 9 and Microsoft Excel. When only two groups were compared, a *t*-test was used. When more than two groups were compared, a one-way ANOVA was performed using multiple comparisons with a Tukey correction. * P<0.05; ** P<0.01; *** P<0.001; **** P<0.0001

## Acknowledgements

We thank the SUNY Upstate Medical University Department of Laboratory Animal Resources for providing care to our mouse colony and the Flow Cytometry Core for access to the flow cytometers. We would like to thank Erica Golemis for the generous gift of the CasL^-/-^ mice. We also thank the members of the Roy laboratory for the discussions, moral support, and critical reading of the manuscript.

## Author Contributions

Conceptualization: L.A.K, N.H.R; Methodology: L.A.K, H.E.S, R.T, M.S, M.K, N.H.R; Formal Analysis: L.A.K, M.S, N.H.R; Writing – original draft and editing: L.A.K, N.H.R; Supervision: N.H.R

## SUPPLEMENTAL FIGURE LEGENDS

**Supplemental Figure 1.**
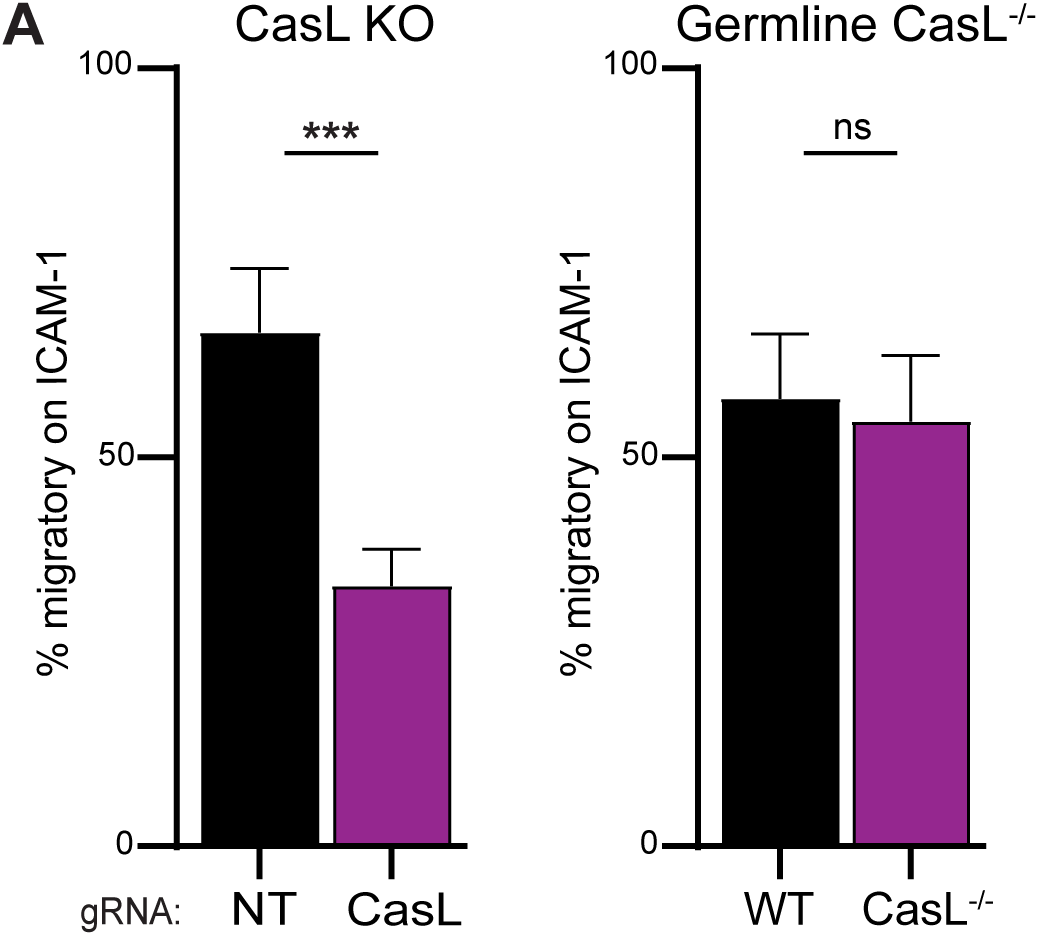
Percent of migratory CD4^+^ T cells on ICAM-1 coated surfaces. Percent migratory CD4^+^ T cells on ICAM-1 using either CRISPR to knockout CasL (n=3, 688 NT cells and 567 CasL KO cells total) or using T cells from germline CasL^-/-^ knockout mice (n=3, 533 WT cells and 612 CasL^-/-^ cells total). T-test, *** = p < 0.001.

**Supplemental Figure 2.**
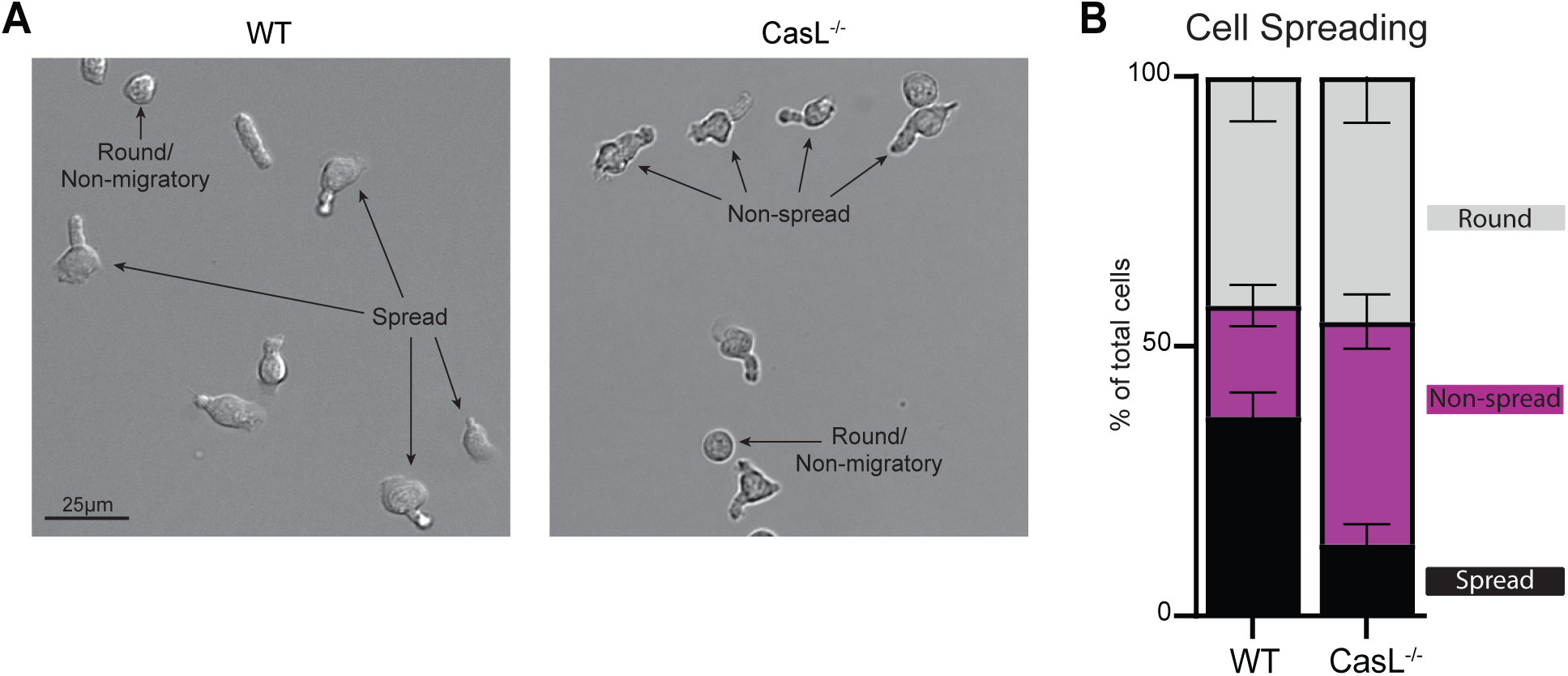
Cell spreading of WT and CasL^-/-^ T cells on ICAM-1. **(A)** Representative images indicating the shape parameter classifications. **(B)** Quantification of cell morphology of WT and CasL^-/-^ CD4^+^ T cells migrating on ICAM-1 (n=3, 248 WT cells and 305 CasL-/- cells total).

**Supplemental Figure 3:**
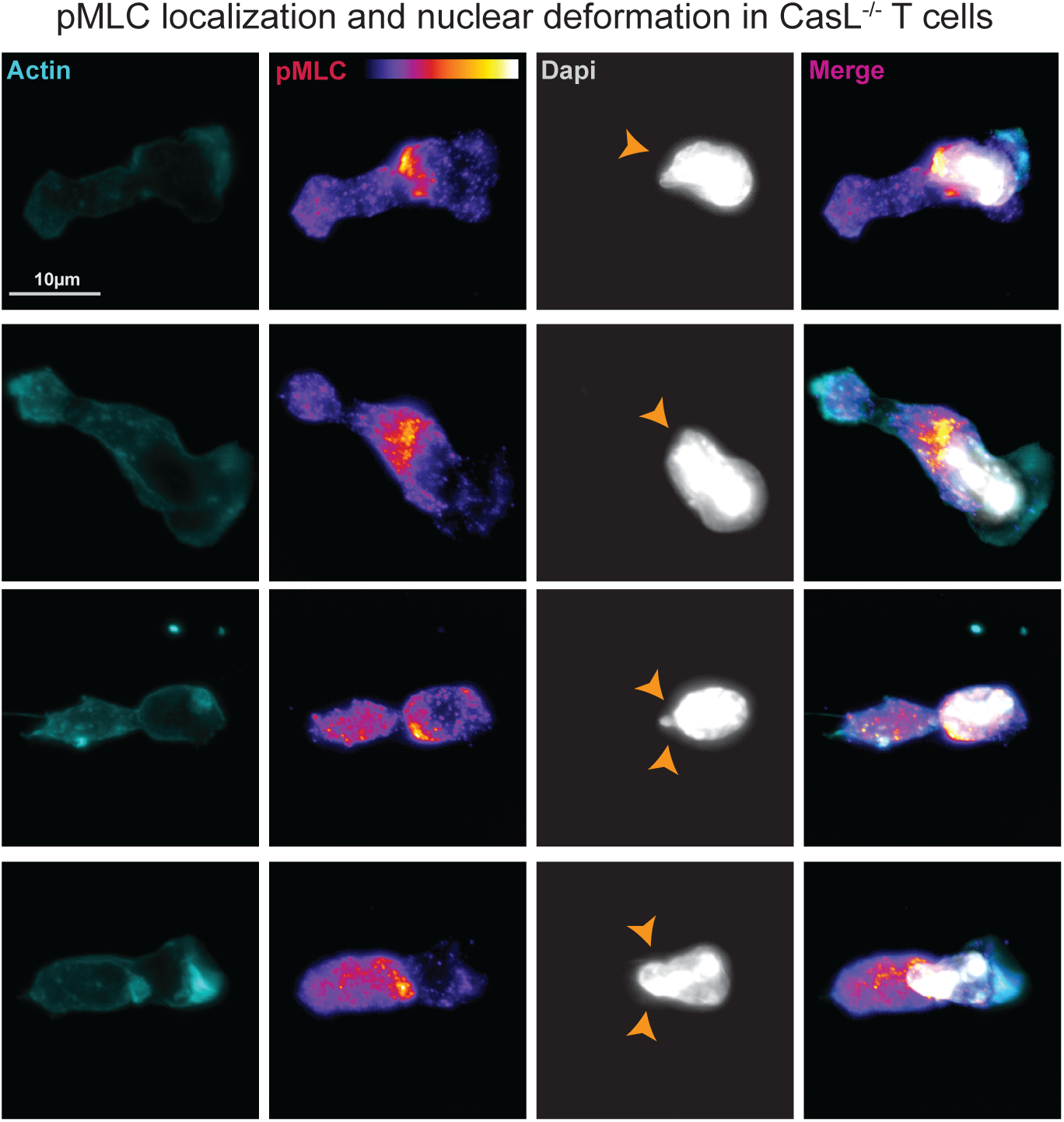
Examples of nuclear deformations in CasL^-/-^ T cells on ICAM-1. CasL^-/-^ T cells were allowed to migrate on ICAM-1 coated surfaces, fixed and stained with anti-pMLC, florescent phalloidin, and DAPI. Darts indicate locations of pMLC and nuclear deformations.

**Supplement Movie 1**: Wild-type CD4^+^ T cell migrating on ICAM-1. 63x, 1sec time points for 1min total.

**Supplement Movie 2**: CasL^-/-^ CD4^+^ T cell migrating on ICAM-1. 63x, 1sec time points for 1min total.

**Supplement Movie 3**: CasL^-/-^ CD4^+^ T cells expressing lifeact-GFP were imaged migrating on ICAM-1. 63x, 1sec time points for 1min total.

**Supplement Movie 4**: WT (non-targeting gRNA) CD4^+^ T cell migrating on ICAM-1 after treatment with 2.5µM Y-27632. 63x, 1sec time points for 1min total.

**Supplement Movie 5**: CasL KO CD4^+^ T cell migrating on ICAM-1 after treatment with 2.5µM Y-27632. 63x, 1sec time points for 1min total.

